# Vitamin B_12_ conveys a protective advantage to phycosphere-associated bacteria at high temperatures

**DOI:** 10.1101/2023.07.30.551168

**Authors:** Margaret Mars Brisbin, Alese Schofield, Matthew McIlvin, Arianna I. Krinos, Harriet Alexander, Mak Saito

## Abstract

Many marine microbes require vitamin B_12_ (cobalamin) but are unable to synthesize it, necessitating reliance on other B_12_-producing microbes. Thus, phytoplankton and bacterioplankton community dynamics can partially depend on the production and release of a limiting resource by members of the same community. We tested the impact of temperature and B_12_ availability on the growth of two bacterial taxa commonly associated with phytoplankton: *Ruegeria pomeroyi*, which produces B_12_ and fulfills the B_12_ requirements of some phytoplankton, and *Alteromonas macleodii*, which does not produce B_12_ but also does not strictly require it for growth. For B_12_-producing *R. pomeroyi*, we further tested how temperature influences B_12_ production and release. Access to B_12_ significantly increased growth rates of both species at the highest temperatures tested (38ºC for *R. pomeroyi*, 40ºC for *A. macleodii*) and *A. macleodii* biomass was significantly reduced when grown at high temperatures without B_12_, indicating that B_12_ is protective at high temperatures. Moreover, *R. pomeroyi* produced more B_12_ at warmer temperatures but did not release detectable amounts of B_12_ at any temperature tested. Results imply that increasing temperatures and more frequent marine heatwaves with climate change will influence microbial B_12_ dynamics and could interrupt symbiotic resource sharing.

## Main Text

Vitamin B_12_ (cobalamin) is required by many marine bacteria and unicellular eukaryotes [1, 2] but is scarce throughout broad regions of the global ocean, forcing microbes that cannot synthesize B_12_ to rely on others that can [3, 4]. Many phytoplankton fulfill their B_12_ requirements through interactions with B_12_-producing bacteria in the phycosphere [5, 6]. Some phycosphere bacteria, like *Ruegeria pomeroyi*, are known B_12_ producers and require B_12_ for growth [6]. Other phycosphere inhabitants, like *Alteromonas macleodii*, cannot produce B_12_ and do not strictly require it for growth but benefit from its availability [7], potentially competing with phytoplankton for B_12_ as has been demonstrated for nitrate [8]. Climate-change-induced temperature increases will influence bacterial growth rates in the oceans [9], but it is unclear how temperature will impact B_12_ quotas and dynamics or downstream effects on microbial communities and interactions. We investigated how temperature stress interacts with B_12_ limitation in phycosphere residents with flexible (*A. macleodii* MIT1002) and absolute (*R. pomeroyi* DSS-3) B_12_ requirements and how temperature stress impacts production and release of B_12_ by a B_12_-producer (*R. pomeroyi*).

To determine the interaction effect of temperature and B_12_ availability on growth, *A. macleodii* and *R. pomeroyi* were grown in a minimal media prepared with (replete) and without (–B_12_) B_12_ across a range of temperatures from 15ºC to 40ºC (Supplementary Information; SI Table 1, SI Figure 1). Lack of exogenous B_12_ significantly diminished *A. macleodii* growth at all temperatures, with the largest effect at the highest temperature (Figure 1). *A. macleodii* biomass was reduced by 57% when grown without B_12_ at the highest temperature in trial 1 (Figure 1B, Supplemental Figure 2), and by 22% in trial 2 (Figure 1A, B). Withholding B_12_ also significantly decreased *A. macleodii’*s mean maximum growth rate (µ_max_; Trial 2): µ_max_ decreased by 0.32 at the highest temperature (27%; *p*<0.05), by 0.14 at the mid temperature (14%; *p*<0.05), and by 0.13 at the cool temperature (18%; *p*<0.05) (Figure 1C). Cell size was largely stable across treatments, but a significant increase was observed at 24 hours for cells grown without B_12_ at the highest temperature in both trials (SI Figures, 6, 7), which is consistent with a reduced growth rate [10] or an arrested cell cycle [11].

**Figure 1.**
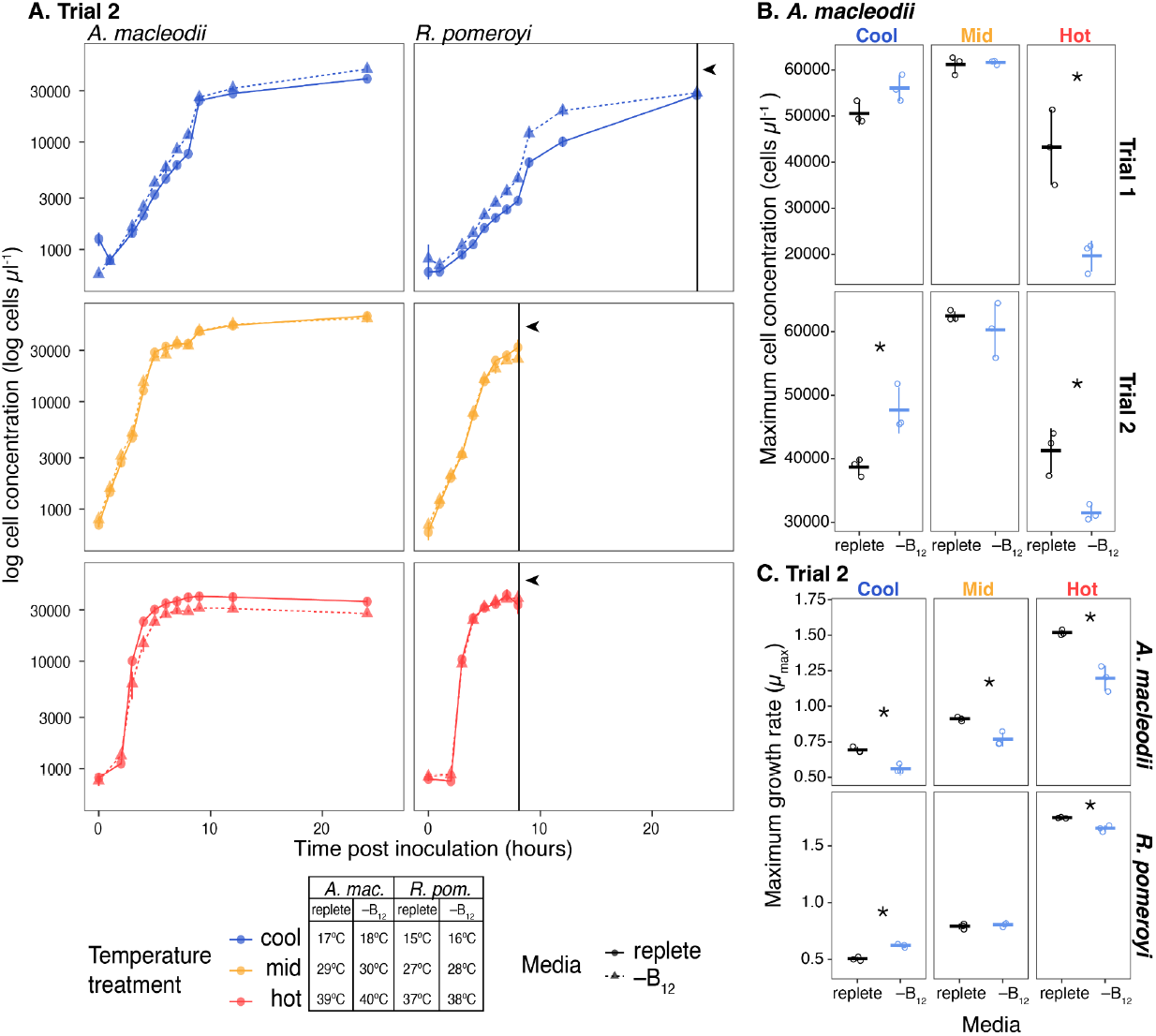
Growth parameters for *Alteromonas macleodii* and *Ruegeria pomeroyi* grown in replete minimal media and minimal media without a vitamin B_12_ source across a range of temperature treatments. (A) Growth curves for both species from experimental trial 2. Colors represent temperature treatments, with the exact temperature for each treatment included in the legend. Point and line shapes represent the media treatment: replete (replete minimal media; circles and solid lines) and –B_12_ (minimal media without vitamin –B_12_; triangles and dashed lines). Each point is the mean log cell concentration of three biological replicates determined by flow cytometry, with error bars representing one standard deviation of the mean. Black vertical lines indicated by arrowheads designate time points where *R. pomeroyi* cultures were harvested for B_12_ measurements by mass spectrometry. (B) Maximum cell concentrations (biomass) reached by *A. macleodii* in experimental trials 1 and 2. Horizontal marks represent the mean cell concentration for each treatment; vertical error bars are one standard deviation of the mean; open circles are individual data points. The statistical significance of media treatment at each temperature was tested by t-test and *p* < 0.05 is indicated on the plots by an asterisk (‘*’). There was a statistically significant reduction in maximum biomass by 57% and 22% in trials 1 and 2, respectively, when *A. macleodii* was grown without vitamin B_12_ at the hottest temperature tested. (C) Maximum growth rates (µ_max_) for *A. macleodii* and *R. pomeroyi* in each temperature and media treatment combination in experimental trial 2. Growth rates were calculated from individual growth curves using the ‘growthrates’ package in the R computing environment. The statistical significance of media treatment on mean maximum growth rate at each temperature was tested by t-test and *p* < 0.05 is indicated on the plots by an asterisk (‘*’). *A. macleodii* cultures grown in replete media had a significantly higher maximum growth rate at all temperatures but the difference in mean maximum growth rate (µ_max_) between media treatments was largest in the hot temperature treatment (0.32 (27%), compared to 0.14 (14%) in mid and 0.13 (18%) in cool). The impact of media treatment on maximum growth rate was more varied for *R. pomeroyi* with the maximum growth rate significantly higher in replete media only at the highest temperature treatment.

**Figure 2.**
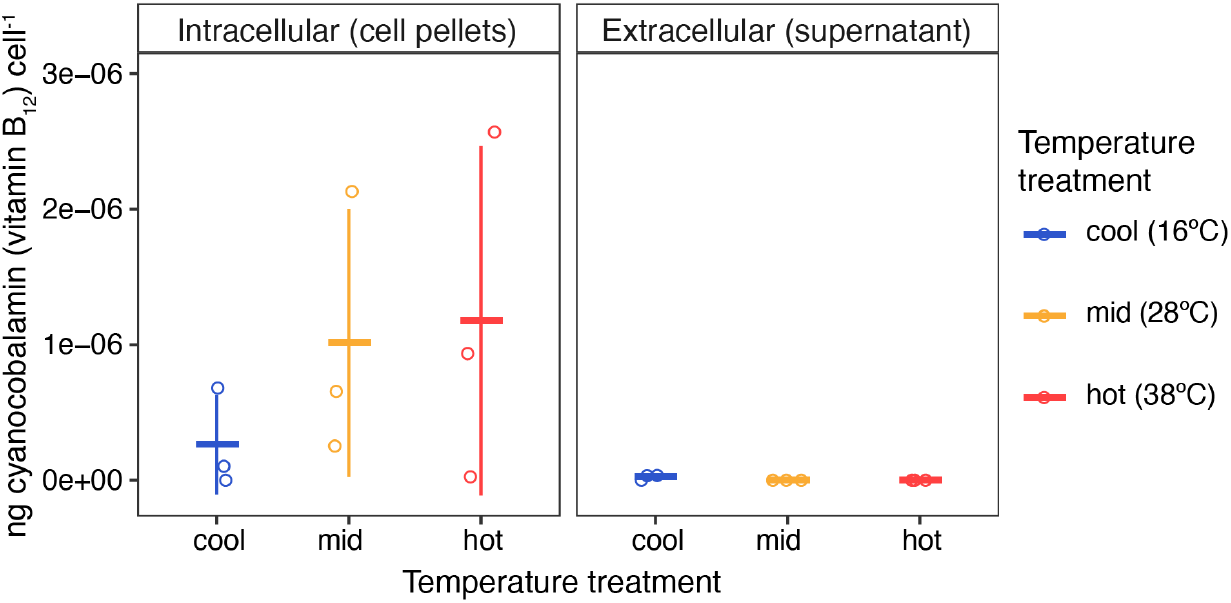
Concentrations of intracellular and extracellular vitamin B_12_ normalized to cell counts in *Ruegeria pomeroyi* cultures grown without an exogenous B_12_ source across three temperature treatments. *R. pomeroyi* cultures in early stationary phase were harvested for cyanocobalamin (B_12_) measurements by mass spectrometry. Measured values were normalized to the number of cells in the originating culture volume (i.e., the number of cells in a cell pellet or the number of cells removed from a supernatant). Horizontal marks represent the mean B_12_ concentration per cell for each treatment; vertical error bars are one standard deviation of the mean; open circles are individual data points. Pelleted cells contained significantly more B_12_ than was present in supernatants (*p* < 0.05, t-test). While not a statistically significant difference, cells grown in the mid and hot-temperature treatments tended to have higher intracellular vitamin B_12_ concentrations than cells grown in the cool-temperature treatment.

The observed changes in growth parameters suggest that B_12_ has a protective or growth-promoting effect in *A. macleodii* at high temperatures. While such observations have not been reported in prokaryotes, B_12_ is protective at high temperatures in the model unicellular eukaryotic algae, *Chlamydomonas reinhardtii* [12]. Like *A. macleodii*, the *C. reinhardtii* genome encodes B_12_-independent (MetE) and B12-dependent (MetH) methionine synthases, meaning it can grow with and without B_12_ [13]. However, exposing *C. reinhardtii* to high temperatures (39ºC) triggers heat shock, chlorosis, and death if B_12_ is unavailable [12]. If B_12_ is available, *C. reinhardtii* exhibits enhanced thermal tolerance, maintaining growth at 42ºC. At high temperatures, *C. reinhardtii* MetE had decreased activity, indicating MetH is more temperature-stable and suggesting a mechanism for thermal protection [12]. This may also hold true for *A. macleodii*. Methionine, however, conveyed a smaller boost in *C. reinhardtii* thermal tolerance than B_12_, advancing the hypothesis that B_12_ enhances thermal tolerance through additional pathways [12]. Notably, B_12_ increases growth in bacteria exposed to other stressors, including oxidative stress [14], low-temperature, and copper stress [15], demonstrating that methionine synthesis at higher temperatures is not the only growth-promoting benefit provided by B_12_ [16].

Exogenous B_12_ had a smaller effect on *R. pomeroyi*’s growth, presumably because it is a B_12_-producer. Withholding B_12_ did not impact the maximum biomass reached by *R. pomeroyi* at any temperature (SI Figure 3) but did significantly decrease growth rates at the highest temperature (Figure 1C). We detected elevated intracellular B_12_ levels in mid and hot temperatures compared to the cool treatment, although not statistically significant (Figure 2). Thus, *R. pomeroyi* may produce more B_12_ at warmer temperatures to maintain similar biomass and growth rates as when exogenous B_12_ is supplied, but B_12_ synthesis cannot keep up with growth requirements at extremely high temperatures. This suggests B_12_ plays a similar growth-promoting or protective role in *R. pomeroyi* as observed for *A. macleodii*. In future studies, this could be tested by growing *R. pomeroyi* mutants incapable of synthesizing B_12_ at high temperatures and determining if growth is diminished when B_12_ is withheld. Of note, extracellular B_12_ was not detected in any of the warm or hot treatment replicates and only trace amounts were detected in two cool treatment replicates (Figure 2). These results imply that little to no B_12_ is released by *R. pomeroyi* in our experimental conditions and that temperature does not have a measurable effect on B_12_ release. While many B_12_-producing bacteria do not release B_12_ [17], these results were surprising because *R. pomeroyi* fulfills the B12 requirement of the diatom *Thalassiosira pseudonana* when grown in co-culture [6]. While co-culture with *T. pseudonana* does not influence *R. pomeroyi* expression of the B_12_ biosynthetic pathway [6], our study suggests that a cue from symbiotic phytoplankton may be required for *R. pomeroyi* to release B_12_.

This study demonstrates that B_12_ conveys a protective or growth-promoting effect at high temperatures for two bacterial species commonly associated with phytoplankton. While the highest temperatures in the study are rare in the current global ocean, they are found in tide pools in subtropical and tropical regions [18], and summer sea surface temperatures (SST) in the Persian Gulf regularly exceed 37ºC [19]. Marine heatwaves—such as the 2023 heatwave affecting the Florida Keys, the Bahamas, and Cuba that caused SST to reach 38ºC (ndbc.noaa.gov)–are expected to become more frequent and severe due to climate change [20]. Our results suggest that increasing temperatures will increase the biochemical need for B_12_ among marine microbial consortia. Shifting B_12_ dynamics may impact symbiotic relationships that sustain phytoplankton and other organisms. Future work should investigate protective mechanisms for B_12_ in marine microbes and the impact of inter-species interactions on B_12_ production and release with changing temperatures.

## Acknowledgments

MMB is supported by a Simons Foundation Postdoctoral Fellowship in Marine Microbiology (award 874439). HA is supported by a Simons Foundation Early Career Investigator in Aquatic Microbial Ecology and Evolution Award (award 931886). AS was supported by the Community College Research Experiences in Woods Hole program (CC-CREW; NSF award ICER-2023192). MAS and MRM were supported by the Simons Foundation, NIH award GM135709-01A1, and NSF award 1850719. AIK was supported by the U.S. Department of Energy, Office of Science, Office of Advanced Scientific Computing Research, Department of Energy Computational Science Graduate Fellowship under Award Number DE-SC0020347. Support for this project also came from the NSF Center for Chemical Currencies of a Microbial Planet (C-CoMP NSF-STC 2019589). We thank Julie Huber and Gretta Serres for organizing and facilitating the CC-CREW program. We thank Erin McParland and Liz Kujawinski for providing the bacterial strains used in this study.

## Data Availability Statement

The raw flow cytometry data generated for this project are publicly available from https://doi.org/10.5281/zenodo.8133026. Vitamin B_12_ mass spectrometry data, intermediate data products, and code used for this study are available in the GitHub repository https://github.com/maggimars/bactB12. The full analysis pipeline is further available as an interactive document: https://maggimars.github.io/bactB12/Flow_Cytometry_Analysis.html.

