## Supplementary Information for "Vitamin B_12_ conveys a protective advantage to phycosphere-associated bacteria at high temperatures"

#### Bacterial Culturing and Flow Cytometry Methods

To determine the interaction effect of temperature and $B_{12}$ availability on the growth of *Alteromonas macleodii* MIT1002 and *Ruegeria pomeroyi* DSS-3, both strains were grown in the cobalt-containing ProMM media (Berube et al. 2015) (SI Table 1), prepared with (replete) and without (–$B_{12}$) vitamin $B_{12}$, across a range of temperatures. Since both strains are able to grow without $B_{12}$, cultures used to inoculate experimental trials were maintained in ProMM media –$B_{12}$ at 19ºC for at least three transfers prior to commencing experimental trials, which prevented contamination of –$B_{12}$ treatment groups with exogenous $B_{12}$ during inoculation. Each treatment condition was inoculated with 100 µl of the same maintenance culture of either strain, which was last transferred 24 hours prior to inoculation. Treatment conditions consisted of 35 ml of sterile media (either replete or –$B_{12}$) in acid-cleaned and autoclave sterilized 50 ml glass culture vials arranged in an aluminum temperature gradient block so that three biological replicates for each strain in each media type were maintained at three temperatures (total $n$ = 36; SI Figure 1).

Growth was monitored in each replicate using flow cytometry following the Eawag protocol for enumerating bacterial cells in water samples (Gatza, Hammes, and Prest 2013). Briefly, 179 µl aliquots from each replicate were stained with SYBR® Green I (Invitrogen) at a final concentration of 1X (i.e., a 10,000X final dilution of DMSO stock solution) in a 96-well plate and incubated for 10 minutes at 35ºC. 50 µl of each sample were analyzed with a BD Accuri C6+ Flow Cytometer using the Eawag bacterial cell analysis template. Flow cytometry data were further analyzed in the R statistical environment (v. 4.2.1; (R Core Team 2018)) with the ‘flowCore’ (v. 2.8.0; (Hahne et al. 2009), ‘ggcyto’ (v. 1.24.1; (Van et al. 2018)), and ‘growthrates’ (v. 0.8.4; (Petzoldt 2022)) R packages. The flow cytometry data generated for this project are publicly available (https://doi.org/10.5281/zenodo.8133026), as are the full analysis pipeline (https://maggimars.github.io/bactB12/Flow_Cytometry_Analysis.html) and intermediate data products (https://github.com/maggimars/bactB12). Flow cytometry measurements were made 3, 6, 12, 24, and 27 hours post-inoculation in the first experimental trial. This sampling scheme was too coarse to estimate maximum growth rates, as there were not enough data points in the exponential growth phase (SI Figure 2). However, a clear decrease in biomass achieved by *A. macleodii* in –$B_{12}$ media compared to in replete media was observed in the hottest temperature treatment (Figure 1B, SI Figure 2). The experiment was, therefore, repeated with a finer sampling scheme where flow cytometry measurements were made approximately hourly for 9 hours post-inoculation (Trial 2; Figure 1A). Replicates of *R. pomeroyi* in the coolest temperature treatment continued to be monitored at 12 and 24 hours post-inoculation since those replicates had less biomass than replicates in the warm and hot treatments and did not appear to have yet reached the stationary growth phase (Figure 1A). All *A. macleodii* replicates were also monitored at 12 and 24 hours post-inoculation.

Growth curves for all replicates in the second trial were analyzed and plotted with the ‘growthrates’ package in the R computing environment (SI Figures 4, 5). The region of the curve with the highest slope was automatically selected for each replicate with the ‘all_easylinear()’

function from the ‘growthrates’ package and the maximum growth rate ($\mu_{max}$) for each sample was exported from the results object. Maximum growth rates were averaged (mean) and the differences between group means were tested with the ‘t.test()’ function in the base R ‘stats’ package with a statistical significance cut-off of $p$=0.05.

**Vitamin $B_{12}$ Mass Spectrometry Methods**

In order to determine if temperature influenced $B_{12}$ production or release by *R. pomeroyi*, replicates in –$B_{12}$ media were harvested in early stationary phase to measure $B_{12}$ concentrations in both cells and surrounding media by mass spectrometry (Okbamichael and Sañudo-Wilhelmy 2004; Tabersky 2010). 30 ml of each replicate were transferred to a 50 ml Falcon tube and centrifuged at 3,234 x g for 10 minutes at 4ºC. Supernatants were decanted to clean 50 ml tubes and both supernatants and pellets were immediately frozen and stored at –20ºC until further processing. For $B_{12}$ extraction, cell pellets were thawed over ice and 10 ml of sterile milliQ water was added to each replicate and vortexed to resuspend the pellet. Pellet samples were kept on ice and sonicated twice for 30 seconds at 50 kHz with a 30-second rest between sonication intervals to rupture cells and release their contents. Cellular debris was removed by centrifuging sonicated pellet samples at 3,234 x g for 20 minutes at 4ºC. Supernatants were decanted to clean 50 ml tubes and brought up to 30 ml volume with sterile milliQ water. Pellet and supernatant samples were acidified with 1M HCl to a final pH of 3.2 before solid phase extraction. Solid phase extraction proceeded using Sep-Pak C18 Classic Cartridges (360 mg sorbant, 55-105 µm; Waters) that were primed by sequentially passing 5 ml LC-MS grade water, 5 ml LC-MS grade methanol, and 5 ml LC-MS grade water through columns. Acidified samples were passed through columns using syringe pressure, desalted with 5 ml of water, and eluted with 2 ml of methanol. Samples were dried overnight with vacuum centrifugation and resuspended in 200 µl of 0.1% formic acid and 2% acetonitrile for same-day mass spectrometry analysis.

Concentrations of intra- (pellet samples) and extra- (supernatant samples) cellular $B_{12}$ (cyanocobalamin) were analyzed by LC-MS on a Thermo Fusion mass spectrometer with a Thermo Dionex Ultimate3000 nanoLC. Each sample was concentrated onto a trap column (0.2 x 10 mm ID, 5 µm particle size, 120 Å pore size, C18 Reprosil-Gold, Dr. Maisch GmbH) and rinsed with 100 µL 0.1% formic acid, 2% acetonitrile (ACN), 97.9% water before gradient elution through a reverse phase C18 column (0.1 x 100 mm ID, 3 µm particle size, 120 Å pore size, C18 Reprosil-Gold, Dr. Maisch GmbH) at a flow rate of 500 nL/min. The chromatography consisted of a nonlinear 30 min gradient from 5% to 95% buffer B, where A was 0.1% formic acid in water and B was 0.1% formic acid in ACN (all solvents were Fisher Optima grade). The mass spectrometer monitored MS1 scans from 677.5 m/z to 680.5 m/z at 120K resolution. MS2 scans were performed on the Orbitrap at 30K resolution with an isolation window of 1.6 m/z with precursor targeted mass of 678.2910 m/z (cyanocobalamin). The resulting raw files were analyzed using Skyline (version 21.2) to calculate precursor and product abundances. A four-point standard curve was prepared with cyanocobalamin solution (certified reference material; Supelco) (SI Figure 8). Cyanocobalamin concentrations in each sample were calculated from cyanocobalamin total fragment area (Figure 2).

| | | |
|---|---|---|
| Macronutrients | $NH_4Cl$ | 0.0032 M |
| | $NaH_2PO_4 \bullet H_2O$ | 0.002 M |
| Trace metals | $ZnSO_4 \bullet 7H_2O$ | 0.0154 mM |
| | $CoCl_2 \bullet 6H_2O$ | 0.0248 mM |
| | $MnCl_2 \bullet 4H_2O$ | 0.00138 mM |
| | $Na_2MoO_4 \bullet 2H_2O$ | 0.0413 mM |
| | $Na_2SeO_3$ | 0.0124 mM |
| | $NiCl_2 \bullet 6H_2O$ | 0.0124 mM |
| | $FeCl_3 \bullet 6H_2O$ | 8.44 mM |
| | $Na_2EDTA \bullet 2H_2O$ | 8.55 mM |
| Vitamins | Inositol | 0.09 M |
| | Thiamine HCl | 0.843 M |
| | Cyanocobalamin | 0.000678 M |
| | Biotin | 0.0122 M |
| | Folic Acid | 0.000552 M |
| | P-aminobenzoic acid | 0.0034 M |
| | Niacin | 0.00616 M |
| | Ca d-pantothenate | 0.0119 M |
| | Pyridoxine | 0.00846 M |
| Organic Carbon Sources | lactate | 0.05% |
| | pyruvate | 0.05% |
| | glycerol | 0.05% |
| | acetate | 0.05% |

**SI Table 1. Components of replete ProMM media.** ProMM media was prepared with 35 PSU autoclaved and 0.2-µm-filtered artificial seawater (Instant Ocean ®, Spectrum) and had a final pH of 7.8. ProMM media without vitamin $B_{12}$ ($–B_{12}$) was prepared identically to the replete media except that cyanocobalamin was omitted. All media was prepared following the instructions for making 1L ProMM media in the “ProSynFest Culturing Manual” (https://www.prosynfest2020.com/wp-content/uploads/2022/03/ProSynFest-culturing-manual.pdf ) and the molarities in the table are calculated based on the protocol in the manual.

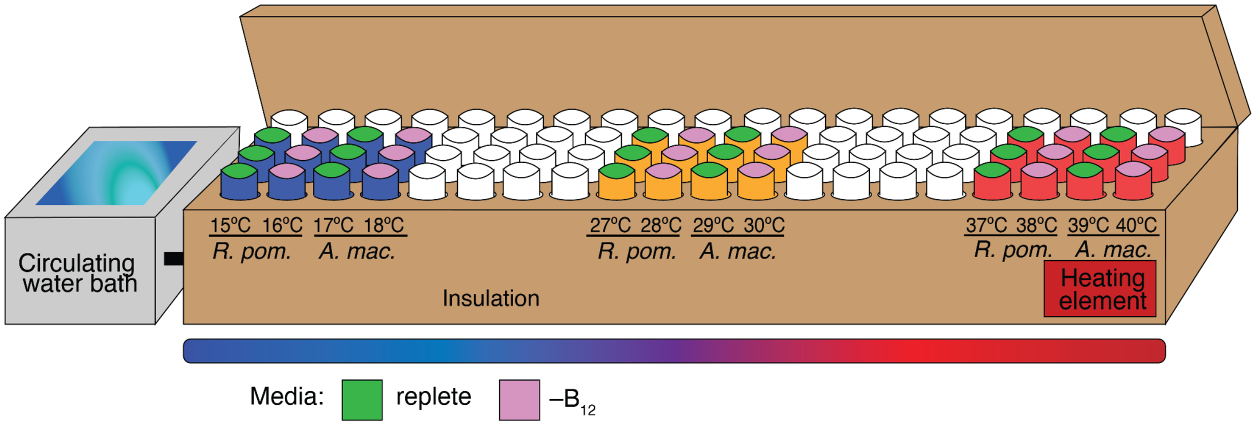


**SI Figure 1. Gradient thermal block experimental design.** The aluminum tube-holding block is situated within an insulated box connected to a circulating water bath at the cold end and a heating element at the warm end. The water bath and heating element temperatures were adjusted to achieve the desired temperature gradient. Blue, yellow, and red coloring of culture tubes in the diagram corresponds to the three temperature treatments in the results figures (cool, mid, and hot). Green and pink coloring of culture tubes in the diagram corresponds to the placement of replete and $–B_{12}$ media treatments in experimental trials.

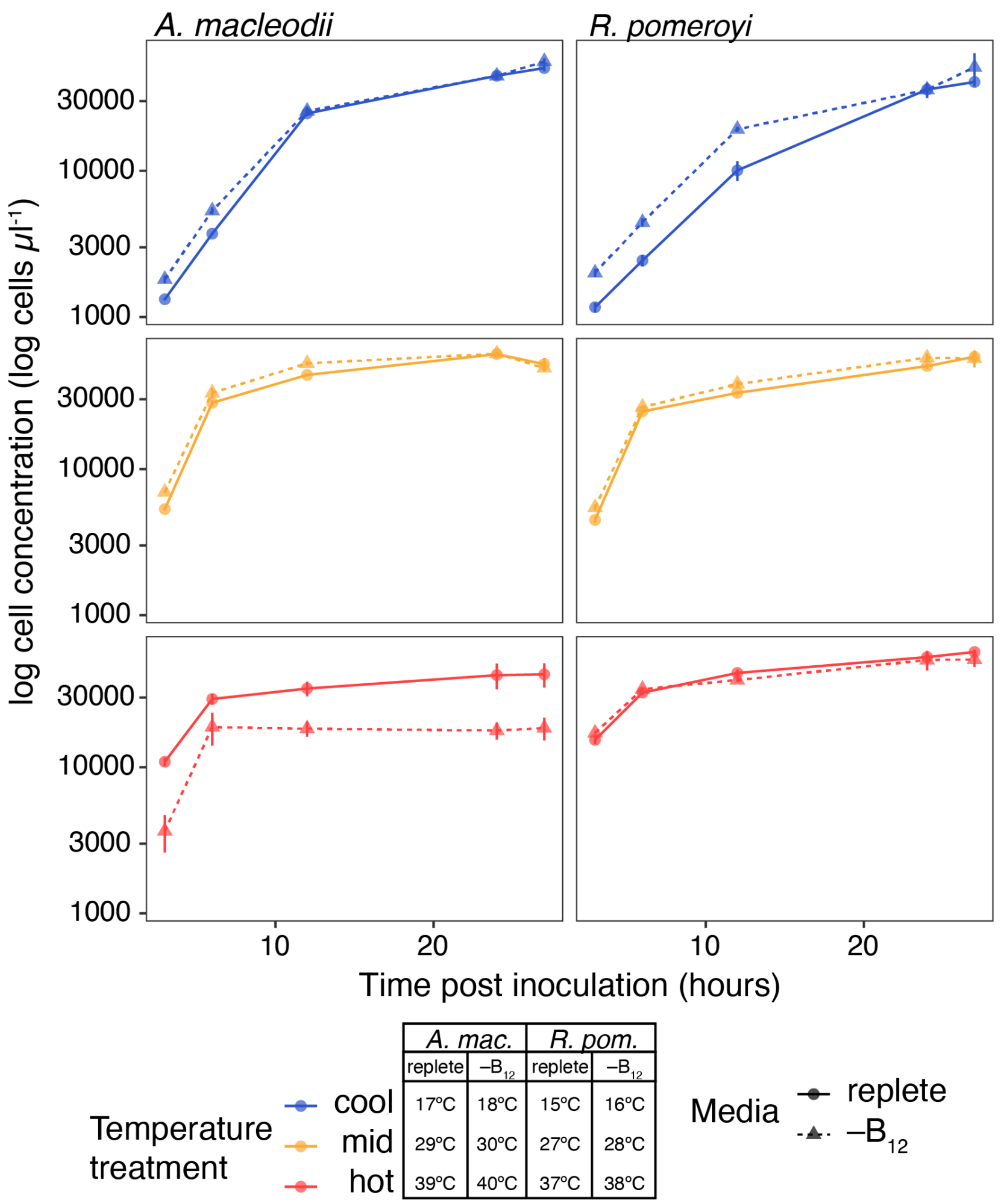


**SI Figure 2.** Cell concentrations through time for *Alteromonas macleodii* and *Ruegeria pomeroyi* when grown in replete ProMM media and ProMM media without a vitamin $B_{12}$ source (–$B_{12}$) across a range of temperature treatments in the first experimental trial. Cell concentrations were measured with flow cytometry in each treatment 3, 6, 12, 24, and 27 hours after inoculation. Colors represent temperature treatments, with the exact temperature for each treatment included in the legend. Point and line shapes represent the media treatment: replete (replete minimal media; circles and solid lines) and –$B_{12}$ (minimal media without vitamin –$B_{12}$; triangles and dashed lines). Each point is the mean of three biological replicates with error bars representing one standard deviation of the mean. The sampling scheme for the first trial did not include enough early samples to evaluate growth rate, however, a reduction in biomass is clearly visible when *A. macleodii* was grown without vitamin $B_{12}$ in the hot temperature treatment.

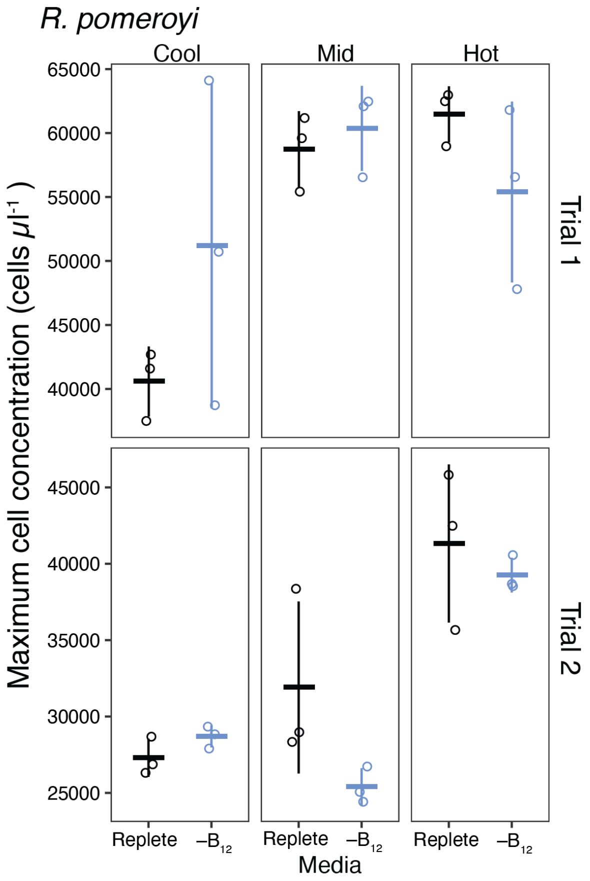


**SI Figure 3.** Maximum *R. pomeroyii* cell concentrations reached in experimental trials 1 and 2. Horizontal marks represent the mean cell concentration for each treatment; vertical error bars are one standard deviation of the mean; open circles are individual data points. The statistical significance of media treatment at each temperature was tested by t-test, but there were no statistically significant differences between treatments.

### *Alteromonas macleodii* (Trial 2)

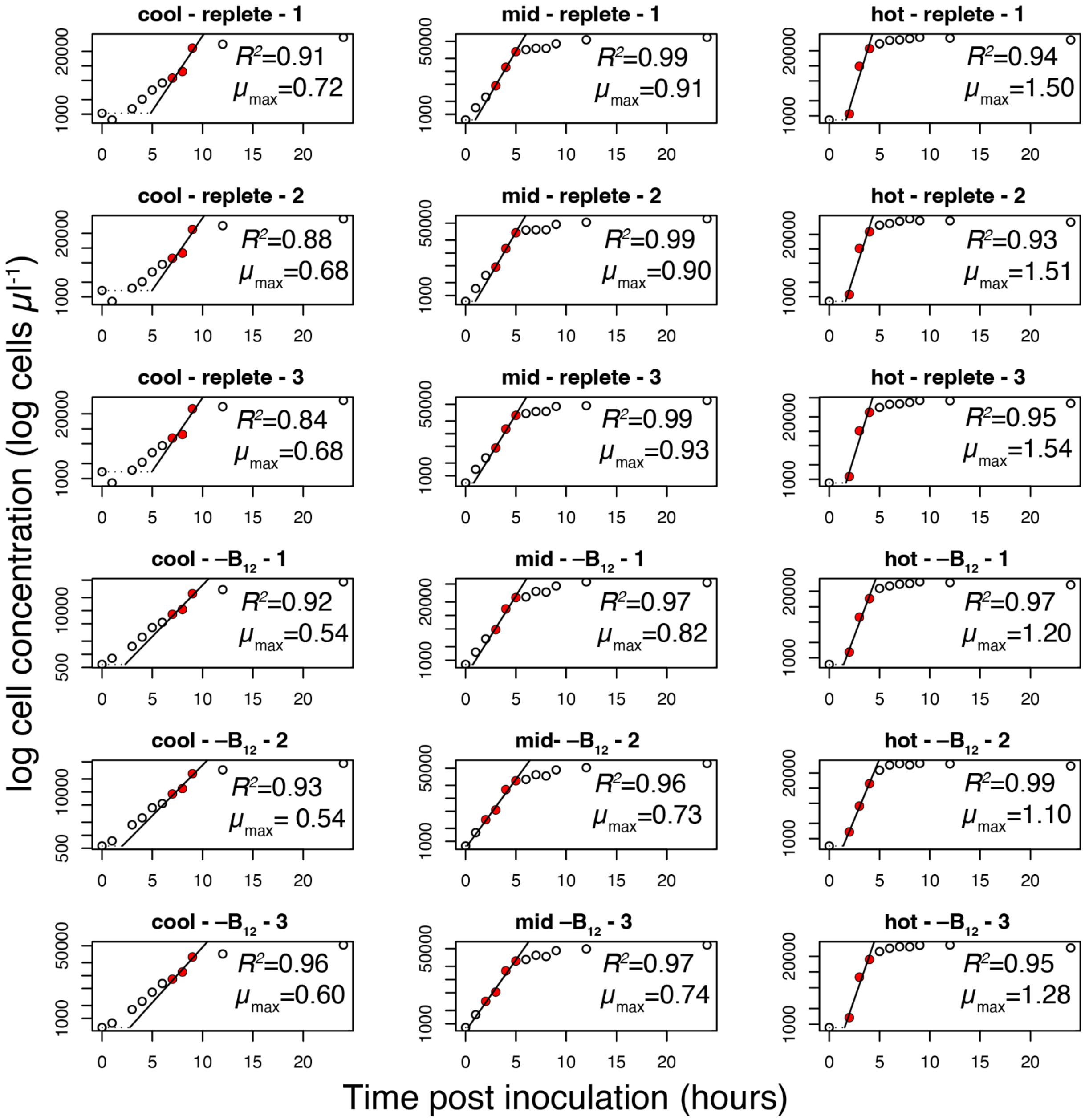


**SI Figure 4.** Individual (per replicate) growth curves for *Alteromonas macleodii* in the second experimental trial. Growth curves were analyzed and plotted with the 'growthrates' package in the R computing environment. The region of the curve with the highest slope was automatically selected for each replicate with the `all_easylinear()` function from the `growthrates` package. The data points in the region of the curve selected for calculating $\mu_{max}$ are filled in red and line fits are plotted as black solid lines. Individual replicate $\mu_{max}$ and $R^2$ are displayed on each plot.

### *Ruegeria pomeroyi* (Trial 2)

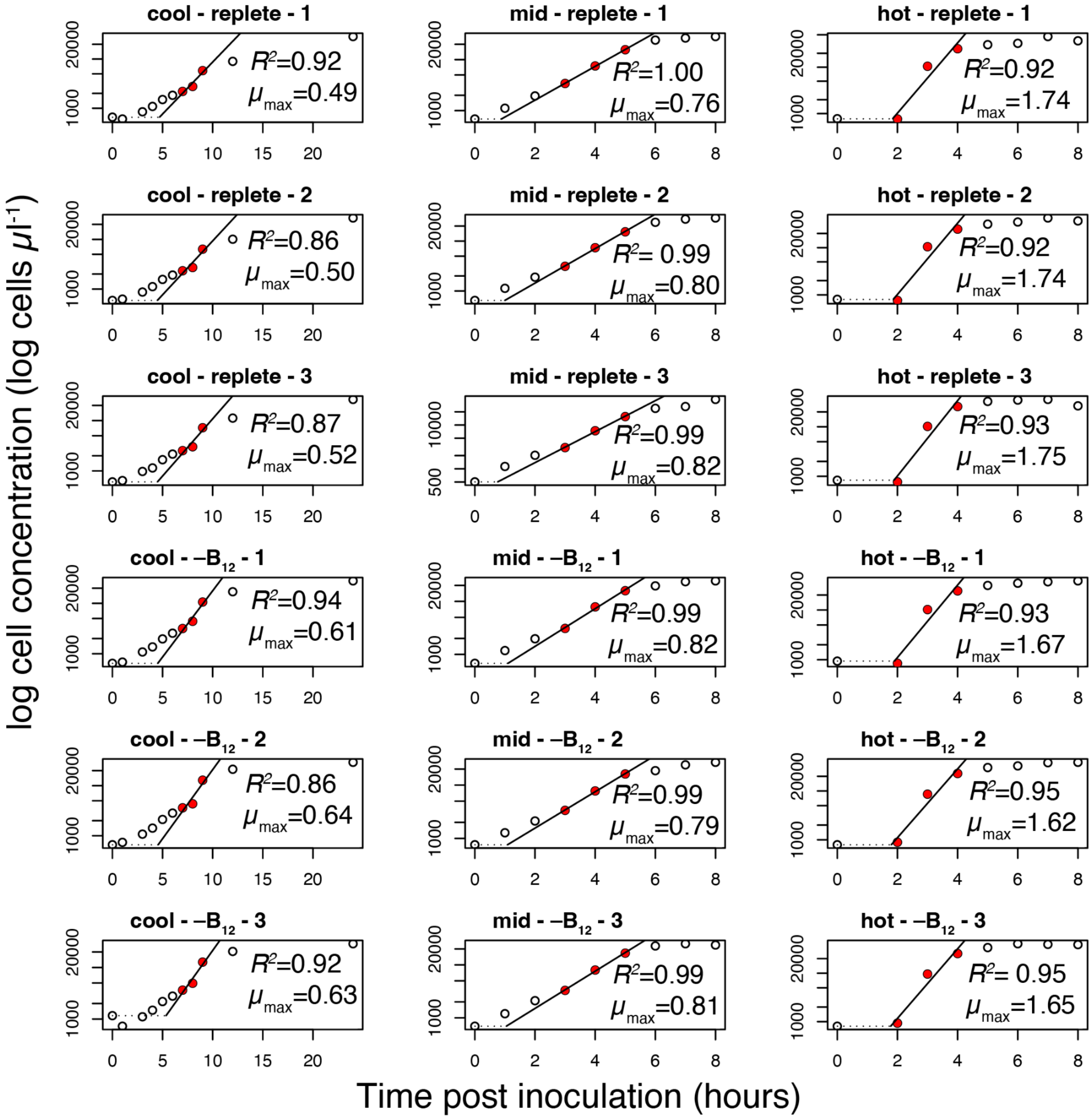


**SI Figure 5.** Individual (per replicate) growth curves for *Ruegeria pomeroyi* in the second experimental trial. Growth curves were analyzed and plotted with the 'growthrates' package in the R computing environment. The region of the curve with the highest slope was automatically selected for each replicate with the \`all_easylinear()\` function from the \`growthrates\` package. The data points in the region of the curve selected for calculating $\mu_{max}$ are filled in red and line fits are plotted as black solid lines. Individual replicate $\mu_{max}$ and $R^2$ are displayed on each plot.

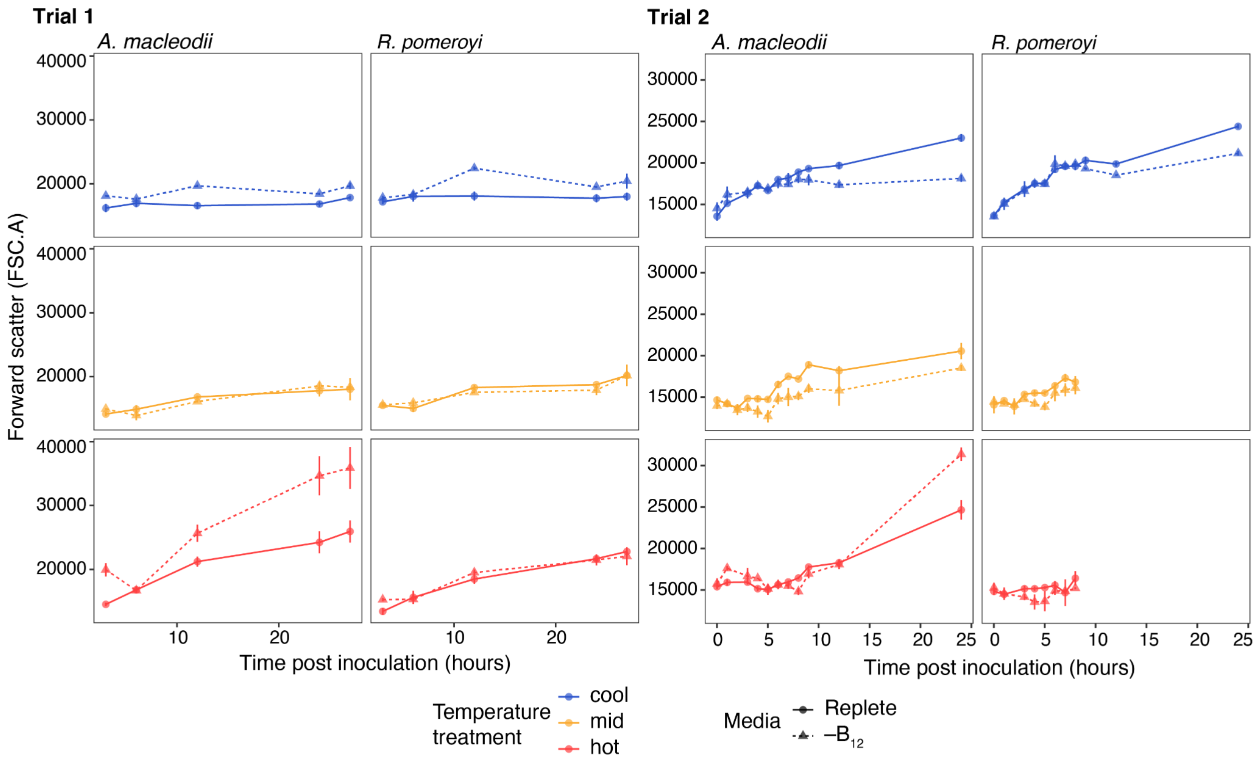


**SI Figure 6.** *A. macleodii* and *R. pomeroyi* cell size through time in experimental trials 1 and 2. Colors represent temperature treatments. Point and line shapes represent the media treatment: replete (replete minimal media; circles and solid lines) and –$B_{12}$ (minimal media without vitamin –$B_{12}$; triangles and dashed lines). Each point is the mean forward scatter of three biological replicates determined by flow cytometry, with error bars representing one standard deviation of the mean. Cell size was largely stable across treatments, but there is a steep increase in *A. macleodii* cell size at 24 hours in the hot temperature treatment in media –$B_{12}$ for both trials.

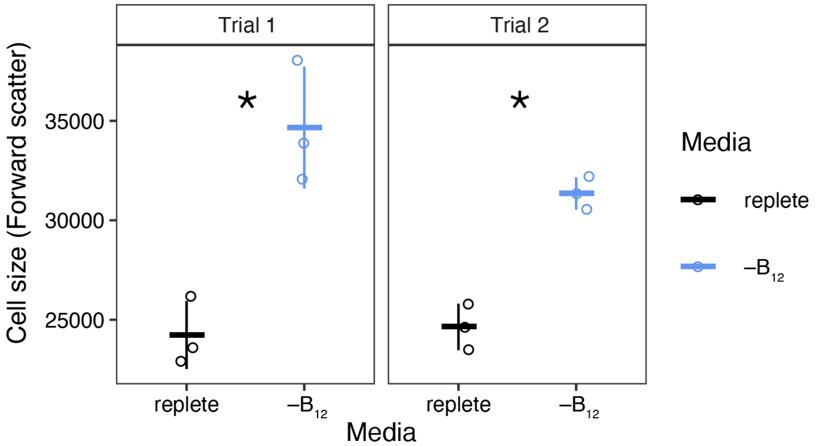


**SI Figure 7.** *A. macleodii* mean cell size (forward scatter; FSC.A) at 24 hours post inoculation in experimental trials 1 and 2. Horizontal marks represent the mean cell size for each treatment; vertical error bars are one standard deviation of the mean; open circles are individual data points (mean cell size per replicate). The statistical significance of media treatment in each trial was tested by t-test and $p < 0.05$ is indicated on the plots by an asterisk ('*'). Cell size was significantly larger in $-B_{12}$ media at 24 hours post-inoculation compared to in replete media in both experimental trials.

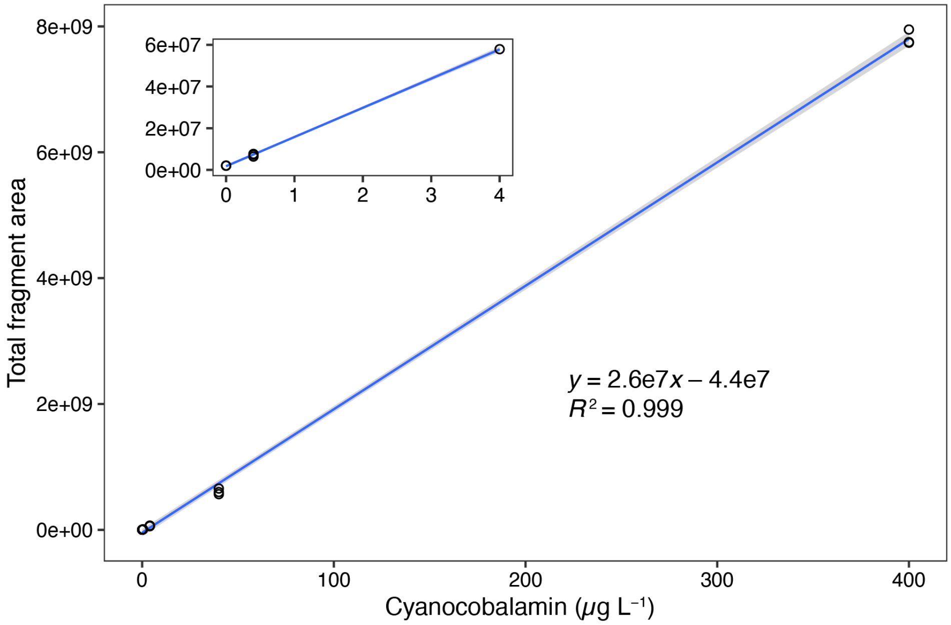


**SI Figure 8.** Calibration curve for quantifying intra- and extracellular $B_{12}$ in *R. pomeroyi* cultures with LC-MS. Measurements were made on 3 replicates each of 0.0, 0.4, 4.0, and 40.0 µg $L^{-1}$ solutions of reference-grade cyanocobalamin. Individual data points are represented by open circles, the blue line represents a linear regression plotted with 'geom_smooth(method="lm"), from the 'ggplot' R package. The inset plot magnifies the lower left portion of the larger plot to show the linear fit of the lower concentration calibration samples. The linear model displayed on the plot was fit with the 'lm()' function in R.

Hahne, Florian, Nolwenn LeMeur, Ryan R. Brinkman, Byron Ellis, Perry Haaland, Deepayan Sarkar, Josef Spidlen, Errol Strain, and Robert Gentleman. 2009. “flowCore: A Bioconductor Package for High Throughput Flow Cytometry.” *BMC Bioinformatics* 10 (April): 106.

Okbamichael, Mussie, and Sergio A. Sañudo-Wilhelmy. 2004. “A New Method for the Determination of Vitamin $B_{12}$ in Seawater.” *Analytica Chimica Acta* 517 (1): 33–38.

R Core Team. 2018. “R: A Language and Environment for Statistical Computing.” https://www.R-project.org/.

Tabersky, Daniel. 2010. “Development and Optimization of a Mass-Spectrometric Method (LC-MS/MS) for the Determination of Vitamin $B_{12}$ in Oligotrophic Ocean Waters.” Edited by U. Telgheder. Master of Science, University of Duisburg-Essen.

Van, Phu, Wenxin Jiang, Raphael Gottardo, and Greg Finak. 2018. “ggCyto: Next Generation Open-Source Visualization Software for Cytometry.” *Bioinformatics* 34 (22): 3951–53.
